# Valence and state-dependent population coding in dopaminergic neurons in the fly mushroom body

**DOI:** 10.1101/809277

**Authors:** K.P. Siju, Vilim Stih, Sophie Aimon, Julijana Gjorgjieva, Ruben Portugues, Ilona C. Grunwald Kadow

## Abstract

Neuromodulation permits flexibility of synapses, neural circuits and ultimately behavior. One neuromodulator, dopamine, has been studied extensively in its role as reward signal during learning and memory across animal species. Newer evidence suggests that dopaminergic neurons (DANs) can modulate sensory perception acutely, thereby allowing an animal to adapt its behavior and decision-making to its internal and behavioral state. In addition, some data indicate that DANs are heterogeneous and convey different types of information as a population. We have investigated DAN population activity and how it could encode relevant information about sensory stimuli and state by taking advantage of the confined anatomy of DANs innervating the mushroom body (MB) of the fly *Drosophila melanogaster*. Using *in vivo* calcium imaging and a custom 3D image registration method, we find that the activity of the population of MB DANs is predictive of the innate valence of an odor as well as the metabolic and mating state of the animal. Furthermore, DAN population activity is strongly correlated with walking or running, consistent with a role of dopamine in conveying behavioral state to the MB. Together our data and analysis suggest that distinct DAN population activities encode innate odor valence, movement and physiological state in a MB-compartment specific manner. We propose that dopamine shapes innate odor perception through combinatorial population coding of sensory valence, physiological and behavioral context.

## Introduction

Behavioral and internal states, past and current experience shape animal perception and behavior. Neuromodulators convey these states and contexts across brain regions and between body and brain [1–3]. Dopamine is among the most intensely studied signals that modulate neural processing and govern plasticity of synaptic connections [4–6]. In the mammalian brain, dopaminergic neurons (DANs) are located in clusters in several brain regions including the mesencephalon, diencephalon and olfactory bulb [7]. The most important sources of dopamine are arguably the *substantia nigra* and the *ventral tegmental area* (VTA) of the basal ganglia, which send projections to the dorsal and ventral striatum, respectively. Brain dopamine has been implicated in cognitive (e.g., motivation, reinforcement, goal-directed behavior, motor control and movement, decision-making, learning) as well as more basic functions (e.g. reproduction, nausea) [4, 7]. How dopamine contributes to these different aspects of neural circuit function and behavior is an open question. A potential answer could lie in the highly localized and region-specific release of dopamine depending on context and task the animal faces [8].

Invertebrates including the fly *Drosophila melanogaster* use dopamine in highly analogous processes [1, 6, 9]. The exquisite tools of fly genetics have provided important insights into the role, molecular and circuit mechanisms of dopamine in associative learning and memory as well as state-dependent behavior (e.g., [10–15]). A focus of many studies has been a dense network of ~200 dopaminergic cells innervating the so-called mushroom body (MB) (Fig. 1A), a brain structure organized in 15 interconnected neuronal compartments (i.e., α1-3, α’1-3, β1-2, β’1-2, γ1-5) [11, 16] (Fig. 1B). These DANs, through unknown sources, respond to stimuli of innate value such as sweetness, heat and electric shock consistent with a model where they convey the ‘unconditioned stimulus’ (US) during learning to MB intrinsic Kenyon cells (KCs) and their corresponding MB output neurons (MBONs) [17]. By taking advantage of highly specific transgenic techniques, recent studies have dissected the function of small subsets or even of single DANs in behavior (e.g., [18–27]). For example, the so-called PPL1 subgroup of DANs, which innervate the α and α’ lobes as well as γ1, γ2 and a region referred to as the peduncle, have been implicated in signaling negative US such as punishment, while PAM DANs projecting to the β, β’ and remaining γ (γ3-5) compartments appear to provide a rewarding contextual signal to KCs and MBONs during associative learning [10, 11] (Fig. 1A).

**Figure 1.**
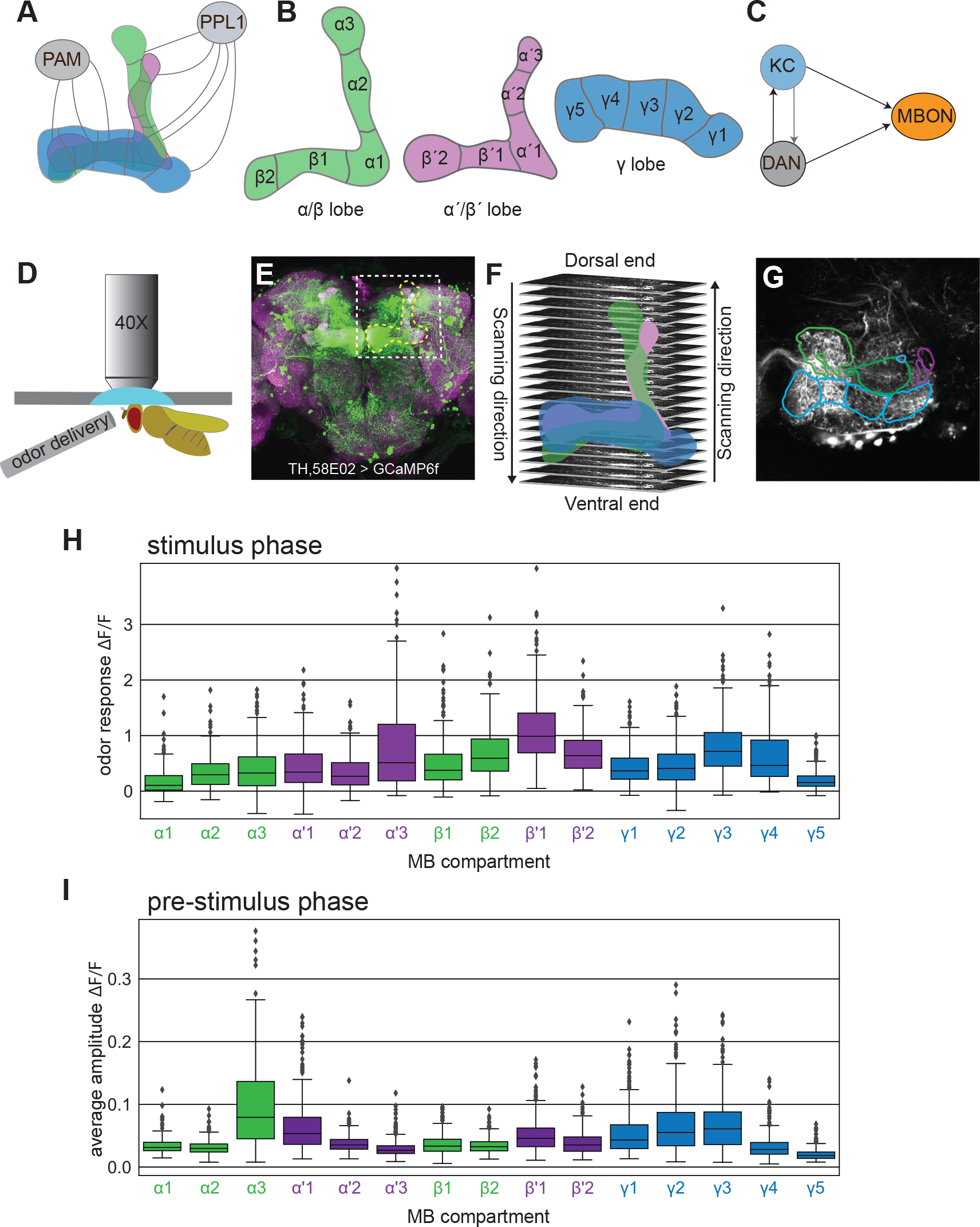
*In vivo* 2-photon population imaging of mushroom body dopaminergic neurons. **(A)** Scheme showing the mushroom body and innervating two clusters of dopaminergic neurons (PAM, PPL1). **(B)** Schematics of the mushroom body lobes with 15 DAN compartments. **(C)** Minimal circuit motif of the mushroom body depicting recurrent connections between dopaminergic cells (DAN) and Kenyon cells (KC). **(D)** A schematic depicting the *in vivo* fly preparation for imaging at the 2-photon microscope. **(E)** A confocal projection image showing the expression pattern of TH,58E02>GCaMP6f. GCaMP6f expression was visualized by anti-GFP (green) and neuropil by anti-discs large (magenta) immunostaining. Dotted yellow line indicates the mushroom body lobes. Dotted white square the region imaged. **(F)** Image showing the imaging planes and scanning directions from dorsal to ventral end covering vertical and horizontal mushroom body lobes. **(G)** An imaging plane showing the 3D masks of the mushroom body lobes used for compartment segmentation and fluorescence extraction. **(H)** Mushroom body compartment-wise odor response. All experiments over all odors, states etc. were pooled. **(I)** Averaged calcium signals during the pre-stimulus phase. Boxes in boxplot show the median and interquartile range, with whiskers showing 1.5 times the interquartile range, and points show outliers.

As yet, less is known about the role of the different DAN types and their respective MB compartments during state-dependent (e.g. physiological states, movement) and innate sensory perception and behavior. Moreover, given that DANs are part of a highly interconnected recurrent network, we lack knowledge regarding their population activity and dynamics as compared to the numerous studies recording the isolated activity of individual DAN types. A notable exception, a study by Cohn et al., provided compelling evidence that relative activities of these neurons matter to US perception and behavior of the fly by analyzing simultaneous calcium signals of all neurons in a DAN subpopulation (~ 40 DANs) innervating the γ-lobe of the MB [28]. This study suggested that subtypes of DANs of different γ-compartments were highly coordinated or anti-correlated in a behavioral state- and experience-dependent manner.

Interestingly, DANs respond to sensory stimuli including odors and temperature changes and contribute to sensory valence decisions in naïve animals [18, 25, 28–32]. Consistently, electron microscopic connectomics data from fly larvae and adults suggests that DANs, especially their axon terminals, receive odor information by KCs as part of a recurrent circuit [33–35] (Fig. 1C). It is also conceivable that olfactory pathways from the lateral horn convey odor information to DANs innervating the MB [36]. These findings motivate the question of whether and how populations of DANs contribute to odor perception and innate valence-decisions, and how they convey state-dependent information. We have begun to address this complex question by recording odor responses across all DANs innervating the fly’s MB. In particular, we analyzed population-wide contributions of DANs to innate valence perception and state changes.

## Results

To this end, we setup an *in vivo* preparation to image calcium fluctuations in all DANs innervating the MB using 2-photon microscopy (*TH-Gal4,58E02-Gal4;UAS-GCaMP6f*; Fig. 1D,E). We recorded from a 210 × 210 pixels area covering the entire MB structure in one hemisphere of the brain, and stimulated every animal with up to two different odorants (e.g. odor 1 when scanning from dorsal to ventral and odor 2 when scanning from ventral to dorsal, Fig. 1F). To minimize bias or experimental artefacts, we randomized odors as well as scanning direction. To compare responses of DAN populations over many individual experiments and animals, we developed a method that allowed us to segment fluorescence changes into each of the 15 MB compartments and to align different brains in 3D (Fig. 1C-E, Fig. S1.2, see Methods). In essence, we used recently-published 3D masks of individual compartments of the MB through a semi-automated landmark registration procedure to assign data from individual pixels to the corresponding compartment [11] (Fig. 1F,G, Fig. S1.2). Importantly, this method enabled us to analyze the responses of the entire 3D volume and not only of a manually defined 2D region of interest for a given compartment. Recording from all DANs at sufficient spatial resolution required imaging the brain over multiple sections with repeated odor presentations of the same odor (see Methods). With these caveats in mind, for this present study, we decided to focus on spatially distinct rather than temporally dynamic signals of MB DANs.

Using this setup, we imaged DAN responses in a total of 165 adult female flies for two out of 12 odorants (i.e. vinegar (Fig. S1A), yeast, citronella, peppermint, 3-octanol, ethanol, 4-methylcyclohexanol, geosmin, isoamyl acetate, 1-Hexanol, 2-Heptanone, 11-cis-Vaccenyl acetate (cVA)) (Fig. S1B). To allow for potential comparison, we chose an overlapping odor set to the one used by Hige et al. [37], who analyzed responses of individual MBONs in naïve animals. Moreover, we recorded from flies of four different internal states, starved (24 and 48 h starved), fed, virgin or mated, to assess the impact of hunger or mating state on DANs. Combining all experiments from all flies, all odor stimuli and states, we first sought to determine how much of the observed variance in the recorded GCaMP fluorescence signal resulted from biological (e.g. odor stimulus, metabolic state, MB compartment) as opposed to technical factors (e.g. imaging direction, order or position of odor stimulus etc.) (Table 1). Using an ANOVA model combining the different factors and their specific compartment effects, we determined that known technical factors only accounted for a small part of the data variance (~1 %). For instance, stimulus order (e.g., whether the odor was the first, second etc. in a series presented to the animal) did not contribute significantly to the observed variability in the data (Table 1). On the other hand, scanning direction (i.e. dorsal to ventral *vs*. ventral to dorsal), and hence number of previously received olfactory stimuli had a significant effect, and explained up to 0.7 % of the variance in the calcium signals (Table 1). This could be indicative of some, albeit mild, adaptation due to repeated odor delivery. The highest contribution (~ 36 %) to the observed variance came, importantly, from biological factors (Table 1). Specifically, differences between the individual MB compartments as well as their response to a specific odorant explained over 30% of the observed variance in the data (Fig. 1H, Table 1). Furthermore, some of the MB lobe compartments showed significant odor stimulation-independent activity during the pre-stimulus phase (Fig. 1I). This was particularly evident for compartments α3, α’1, γ1, γ2, and γ3 (Fig. 1I). Interestingly, these were not the same compartments that showed the highest variance in their odor responses (i.e. α’3, β’1, γ4), implying that these variabilities are a biological feature of the respective compartments, and not a technical artefact that should conceivably affect always the same lobes (Fig. S1C).

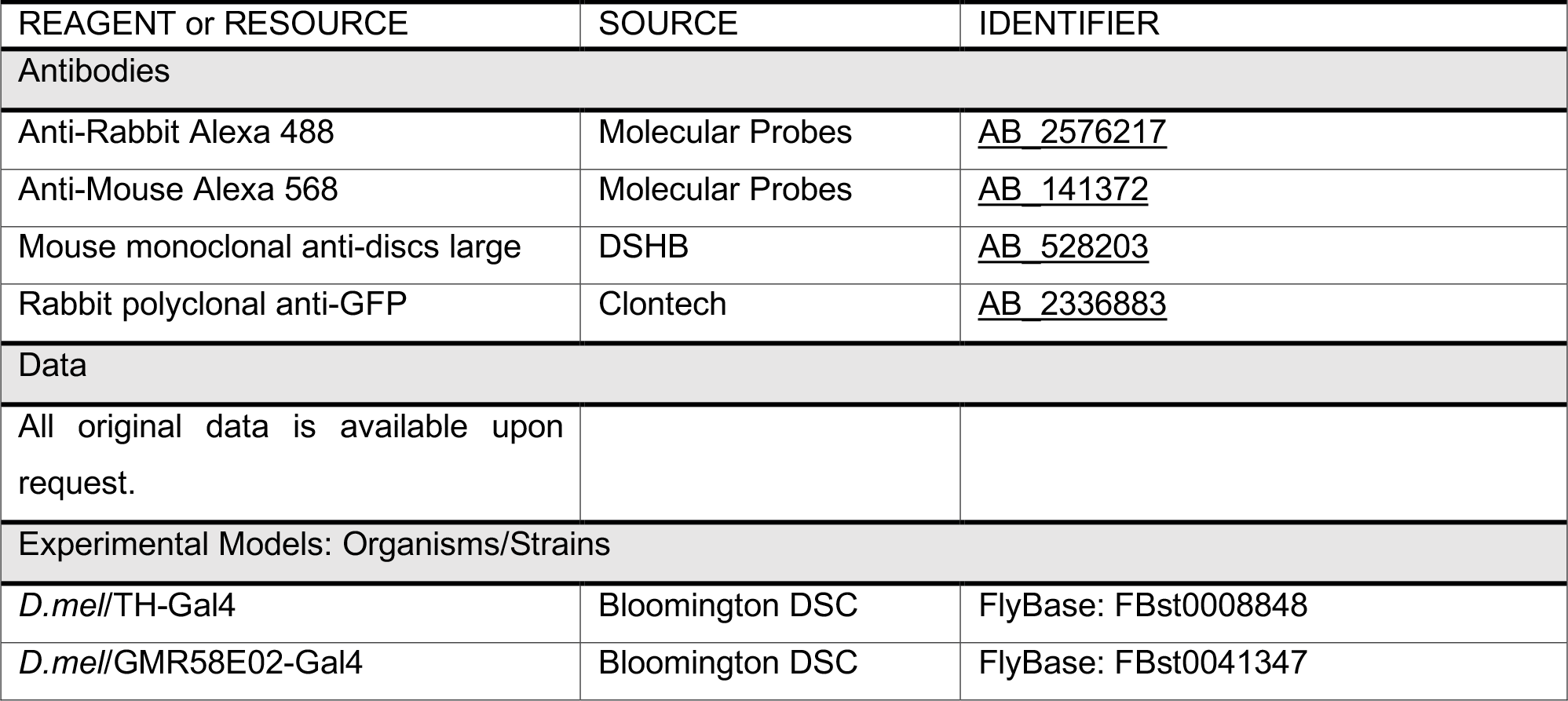
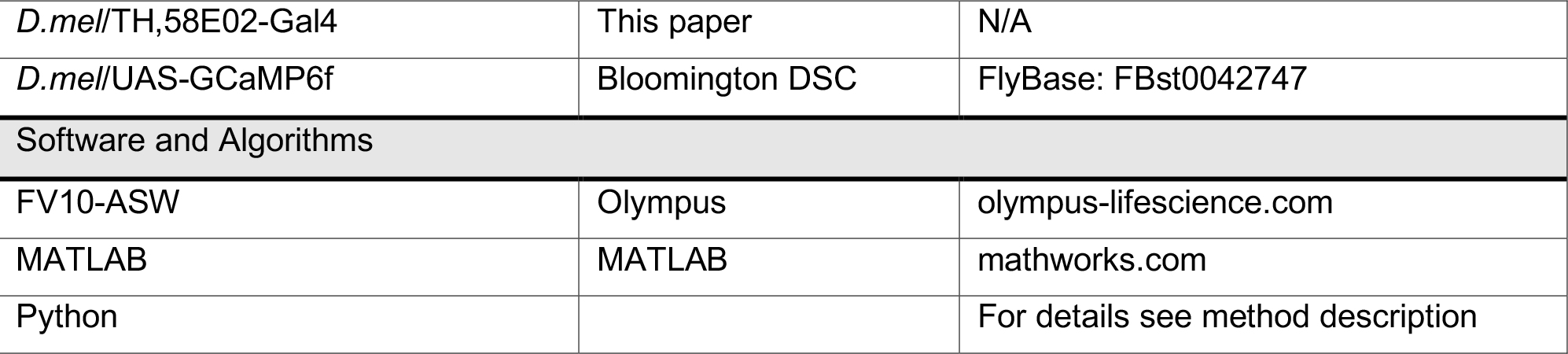
Key Resources Table.

Hence, by considering the known and intended conditions of the experiments (i.e. odor identity, metabolic state, compartment identity, imaging direction, position in odor sequence), we were able to explain ~ 37% of the observed variance in our data (R^2^ = 0.374, Table 1). The highest contribution came from the differences in activity of different MB lobes indicating that distinct MB compartments display characteristic calcium responses (Table 1).

We next asked whether and what type of information DAN responses contain about the odor stimulus and if they might impact how an odor stimulus is perceived by the animal. In order to visually compare responses to different odors, we plotted averaged odor responses over all animals that received a given stimulus in a 2D heatmap (Fig. 2A, S2). Considering these average response heat maps, we wondered whether DANs could help to encode odor identity in their population activity (Fig. 2B). As indicated in Table 1, stimulus identity induced compartment-specific responses, and in total accounted for 12.1 % of the observed variance. Of note, the specific effects of the stimulus on different compartments explained a larger proportion of the variance than just stimulus alone (Table 1, 8.8 % vs. 3.3 %), suggesting that different compartments might respond differentially to different stimuli. Decoding odor identity from the recorded DAN population activity using logistic regression (see Methods) indeed performed at more than three times higher accuracy than chance level (26% vs. 8% chance level) (see confusion matrix in Fig. S2B). Consistent with this, odor representation in Linear Discriminant Analysis (LDA, see Methods) space was not homogeneous, with the distance between each odor cluster higher than if odor had been randomly shuffled (1.65 vs. 0.79 ± 0.06; see for example Fig. 2C, D). Although DANs are not perfect encoders of odor identity in the fly brain, these results suggest that their responses might still convey some potentially useful information about the type of odor that the animal is smelling.

**Figure 2.**
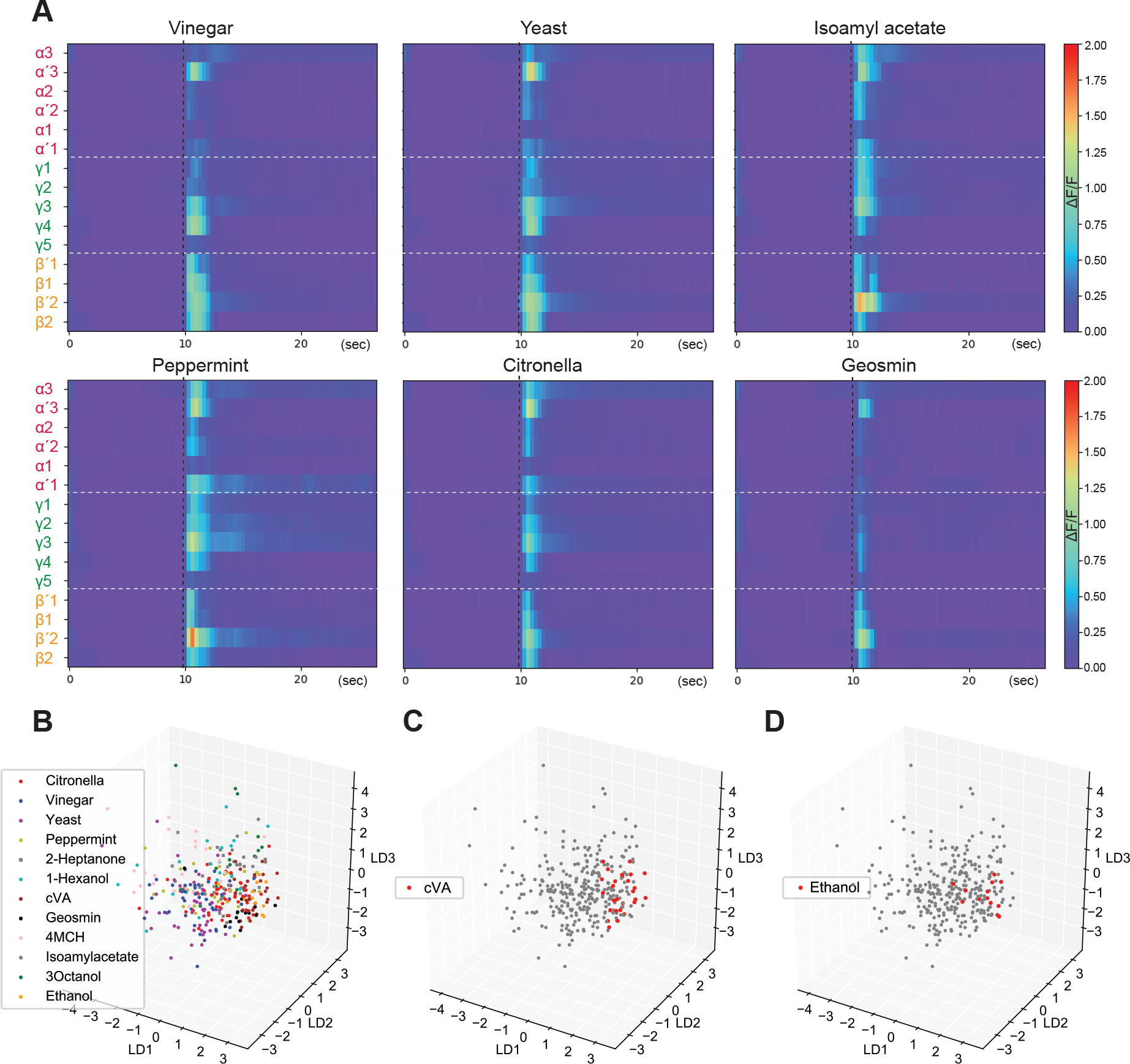
Odor identity, spontaneous activity of distinct lobes, movement. **(A)** Heatmaps showing averaged DAN responses to different odors in 15 mushroom body compartments (n = 319 experiments, 165 female flies, all states). Vertical dashed black line indicates stimulus delivery time. Horizontal dashed white line separates groups of lobes. **(B-D)** State space after linear discriminant analysis dimensionality reduction. **(B)** presents all odors, while **(C)** and **(D)** highlight cVA and ethanol.

Humans frequently rate odors, including novel scents, as pleasant or unpleasant [38]. In flies, DANs have been implicated in regulating innate olfactory preference behavior (e.g., [13, 15, 18, 39]). We therefore next looked for an innate valence code for odors in this neuron population. To this end, we focused on the odors reported to possess an innate valence in behavioral assays for the animal: The food odors vinegar and yeast were categorized as positive and citronella, geosmin, and peppermint as negative. Using the same ANOVA model as above, but analyzing stimulus valence instead of stimulus type, we found that valence accounted for 3.6 % of the observed variance as compared to 12.1 % for stimulus type (Table 1 and 2). This difference of 8.5 % is consistent with the interpretation that DANs, in addition to valence, hold some further information about the nature of an odor stimulus (e.g. odor type).

Regarding odor valence, by examining post hoc pairwise comparisons and regression coefficients for individual compartments, and noticed that DANs innervating the α3, α’1 and γ2 compartments responded significantly stronger to aversive odors as compared to appetitive odors (Fig. 3A). Conversely, β2, β’2 and γ4 showed higher DAN responses for odors of positive valence (Fig. 3A). This division coincides with the PAM / PPL1 MB innervation boundary and their reported responses to stimuli of opposite innate valences such as sugar, bitter tastes or electric shock [20, 27, 29]. Interestingly, using logistic regression on the DAN population activity data, the valence of an odor could indeed be decoded at a much higher confidence than chance level (70%, p = 0.0006), suggesting that odor evoked activity within the DAN network could contribute to an animal’s innate perception of odor valence (Fig. 3B). Moreover, LDA loadings (the weights of the original data points projected onto the LDA direction) were also segregated along opposite odor valences (Fig. 3B; d’ = 1.29 compared to 0.60 ± 0.11 for random assignment), and LDA coefficients showed that DANs innervating different lobes contribute differentially to this segregation. The most significant contribution to odor valence classification was found for DANs projecting into the β’2 compartment (Fig. 3C).

**Figure 3.**
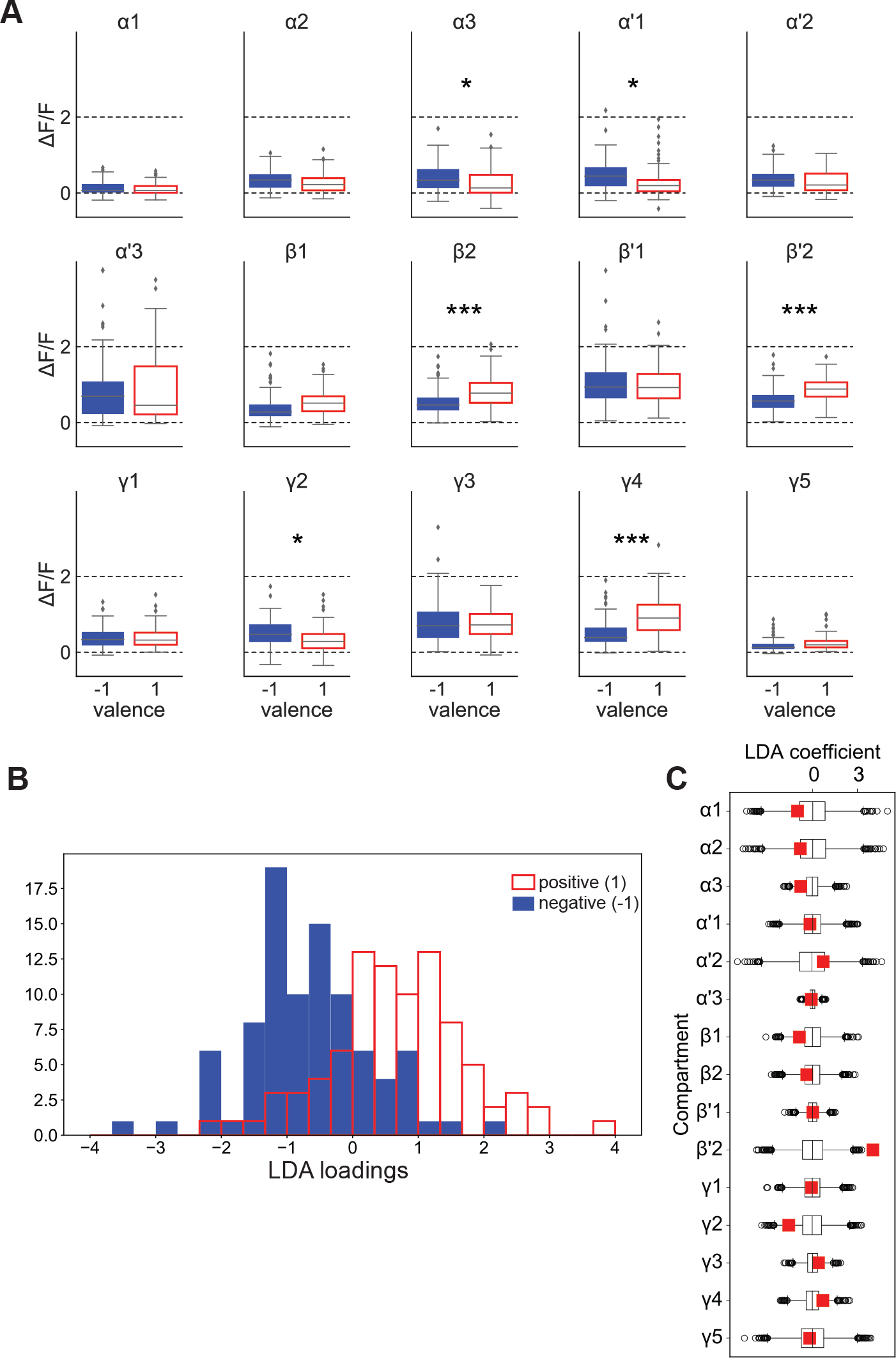
Dopamine neuron responses reveal innate odor valence. **(A)** Lobe-wise pairwise comparison of response to positive or negative valence odors. Stars represent t-test multiple comparison corrected p-values. Stars represent t-test multiple comparison corrected p-values. * p<0.05, ** p<0.01, *** p<0.001. **(B)** Histogram of loadings of projection on the valence optimized LDA dimension. **(C)** Lobe coefficients for valence optimized LDA in color. Box plots represent chance level.

Together, these data provide new insights into how the population of DANs could contribute to innate valence perception of odors as previously observed in behavioral studies. Curiously, the obtained 70% confidence level roughly corresponds to the percentage of animals that show strong innate attraction or aversion to appetitive or aversive odors, respectively, in simple two-choice behavioral assays such as the T-maze [40–42].

Metabolic state is an important determinant in odor perception for many animals including humans [43–45]. We thus compared DAN responses of fed to the responses of 24 and 48 h starved animals (Fig. 4A). Metabolic state indeed contributed significantly, albeit less than stimulus valence, to the variance explained in the ANOVA analysis based on data from all test odors and all recorded animals (0.7 %, Table 1). Both post hoc pairwise comparisons and regression coefficients indicated that not all compartments contributed equally. We detected the most significant differences in responses in compartments α1, β1, β’1, and β’2 (Fig. 4A, S4A). Previous studies that focused on individual DANs implicated some of these compartments in metabolic state-dependent choice behavior (i.e. β’2) [13, 18, 46]. Our present results suggest that other compartments such as β’1 might also contribute in conveying feeding state to sensory processing. Importantly, decoding the responses of the entire DAN population allowed us to predict whether the fly was starved or fed with a 61% confidence level over chance (p = 0.04). LDA projections as done above for odor valence, segregated between starved and fed animals (Fig. 4B; d’ = 0.89 compared to 0.49 ± 0.09 for random assignment). Interestingly, compared to the DAN population activity no single compartment was sufficient to predict starvation level with significant confidence (Fig. 4C). Taken together, the results suggest that feeding state not only modulates the response of DANs to odors, but that the population of DANs are significantly better at conveying the animal’s feeding state to its higher cognitive centers than individual DANs or MB compartments.

**Figure 4.**
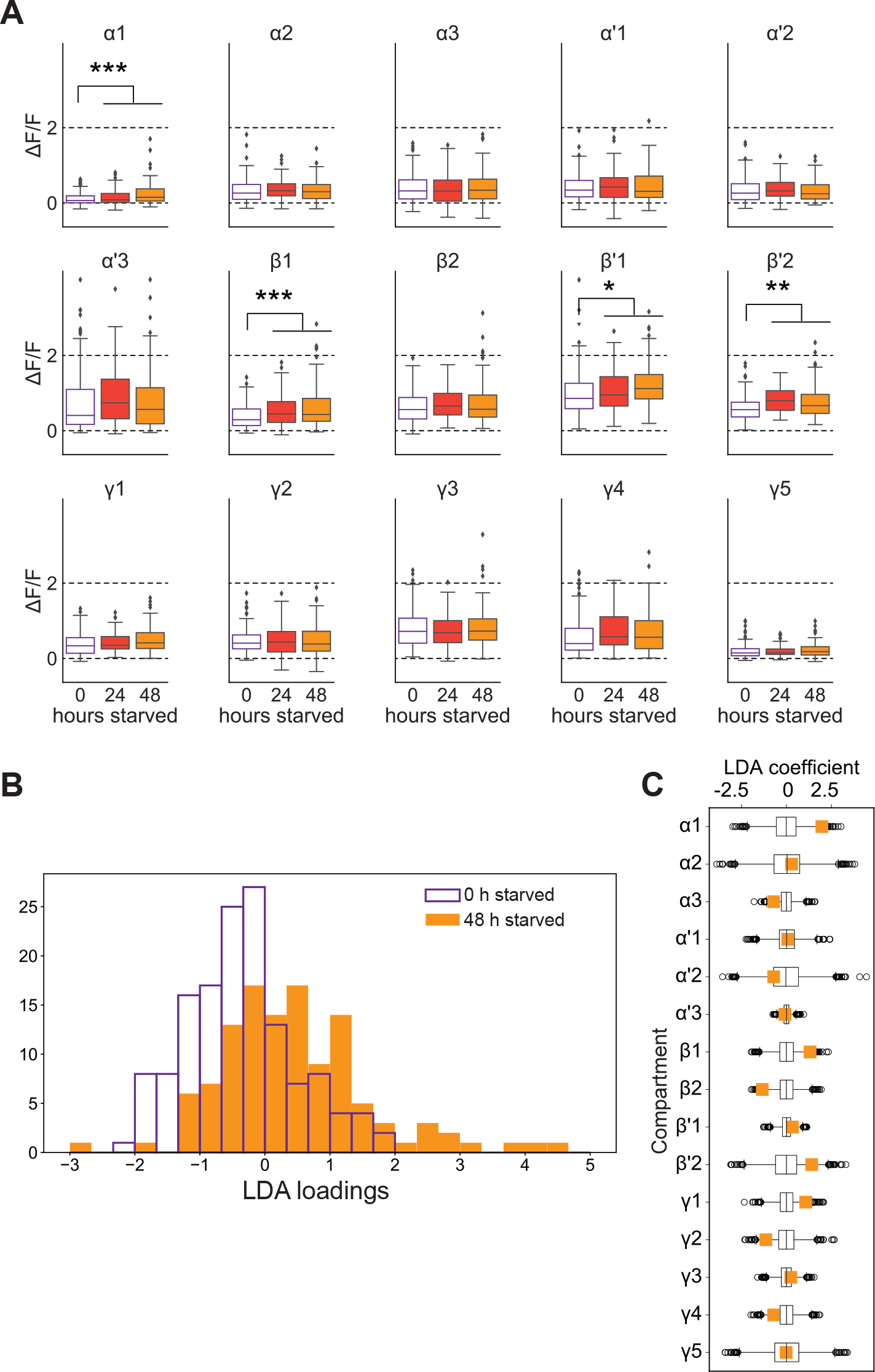
Metabolic state affects dopamine neurons in specific mushroom body compartments. **(A)** Lobe-wise comparison of odor responses in different starvation states. Stars represent t-test (data for 24h and 48h starvation were pooled) multiple comparison corrected p-values. **(B)** Histogram of loadings of projection on the starvation state optimized LDA dimension. Stars represent t-test multiple comparison corrected p-values. * p<0.05, ** p<0.01, *** p<0.001. **(C)** Lobe coefficients for valence optimized LDA in color. Box plots represent chance level.

One of the odors in our set was the sex pheromone cVA. Mating experience changes how males and females perceive of their environments and members of their own species (e.g. [23, 47–49]). Thus, we compared cVA DAN responses in mated females to that of virgins (Fig. S4B,C). In particular, DANs projecting into compartments α3, β’1 responded differentially to cVA and significantly predicted mating state in a regression model (Fig. S4C,D). These results indicate that reproductive state also influences how the DAN population encodes odors.

Given that some DANs were shown to be active during movement of the fly, independent of an external sensory stimulus, we wondered whether and where movement-related activity was encoded by the MB DAN population [28, 50, 51]. To test this, we analyzed another dataset of tethered flies walking on a ball, but which were not stimulated with odor. These data were obtained by using light field microscopy allowing us to image calcium signals induced by movement at a very high temporal resolution across all compartments [51]. We found that some MB compartments were strongly responsive to movement. In particular, regression analysis indicated that calcium signals in compartments β1, β2, β’2 and γ3-5 correlated with walking (Fig. S5A,B). The average regression coefficients for walking ranged from 0.01 – 0.07 (Fig. S5B), which was similar to the coefficients observed for metabolic state (0 – 0.12, Fig. S3), mating (−0.02 – 0.35, Fig. S4D) and valence (−0.09 – 0.22, Fig. S4A). These data imply that movement is encoded by the MB DAN population and might impact on an animal’s sensory perception to a similar degree as other internal states.

## Discussion

Taken together, we find that different DANs within the population of MB-innervating neurons respond differentially to different odors and states, and encode information regarding odor valence and physiological state in a compartment-specific manner (Fig. 5). Different compartments show strong responses when the animal is stimulated with a given odor, whereas others display higher intrinsic variability in the absence of stimuli (Fig. 5). Furthermore, some compartments are clearly modulated while the animal is walking (Fig. 5). Similarly, odor valence is encoded more reliably by certain compartments than others when comparing DAN responses across the entire MB DAN population (Fig. 5). And finally, metabolic and reproductive states modulate odor responses of the DAN population; DANs innervating some compartments again showed significantly higher effects of modulation as compared to others (Fig. 5).

**Figure 5.**
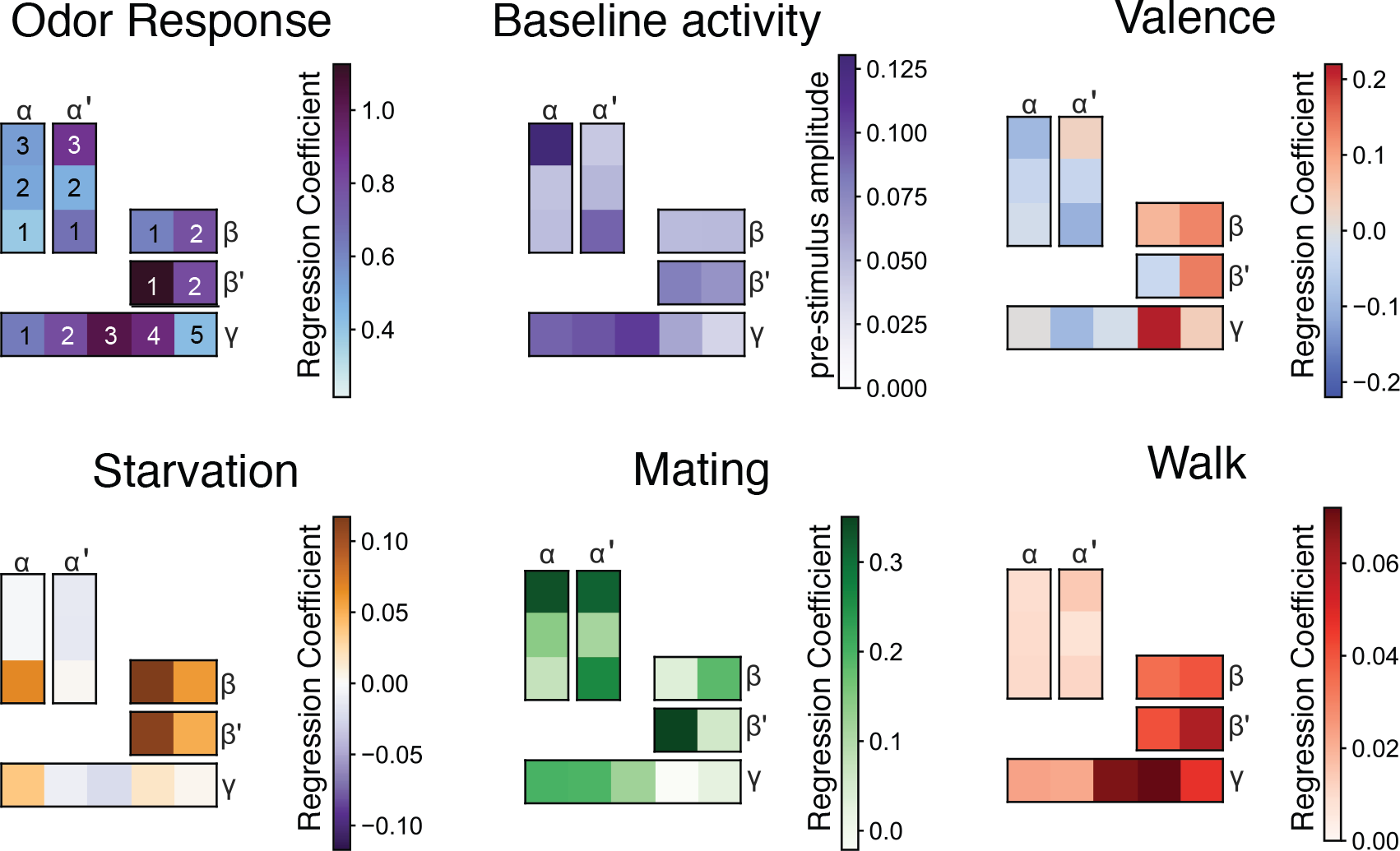
Summary models of the contribution of different MB compartments. Mushroom body compartments are color coded with the value of the regression coefficient or mean amplitude of baseline activity in the absence of an odor stimulus. Different compartments could therefore contribute differentially to the perception of odor valence and internal state.

A putative caveat in our data is the observed large variance that cannot be explained with known or controlled variables (Table 1). This variance, however, could be consistent with the finding that different individual flies respond differently to odors and state changes resulting in the considerable variability typically observed in innate preference behavioral experiments. It is possible that individual experience shapes odor responses as suggested for variance observed in MBONs of different flies [37]. Alternatively, stochastic processes during development could lead to different wiring of neurons in individual flies [52], ultimately shaping how DANs respond to a given odor. Finally, we cannot exclude that differences in GCaMP expression and compartmental localization, as well as differences in fly preparation contribute to the different responses. We hope to answer how individuality shapes odor experience at the level of DANs in the future by combining higher throughput imaging methods with developmental genetics.

Importantly, in spite of putative individual differences, DANs as a population contain important information that could contribute markedly to the innate and state-dependent perception of odors by gauging output pathways of the MB and possibly of the lateral horn. More specifically, compartmentalized but population-wide DAN responses to odor could bias MBON output by acutely modulating KC-MBON synapses. In this scenario, odor information could be received from the LH, but might also originate from KC – DAN synapses directly. In conclusion, we propose that DANs as a population govern innate perception and behavior by directly and differentially encoding the innate valence of a sensory cue and the animal’s current physiological and behavioral state.

## Acknowledgements

We would like to thank Sebastian Onasch and Ariane Böhm for their help with data analysis and modeling, and Christian Schmid for writing the odor delivery script. We are grateful to Rüdiger Klein and the Max Planck Institute for Neurobiology for generously sharing their multiphoton microscope, and to Robert Kasper for his technical help. This study was financed by the German Research Foundation (DFG, SFB870 (A04)) and an ERC starting grant (FlyContext) to IGK.

## Author contributions

K.P.S and I.G.K. conceived the study in collaboration with R.P. and V.S. K.P.S. carried out all *in vivo* two photon imaging experiments. V.S. and R.P. designed the 3D image analysis method. K.P.S. and V.S. extracted the raw data from the two photon experiments and S.A. with help of V.S., R.P. and J.G. analyzed the data and generated the models. S.A. carried out all light field microscopy experiments and their analysis. I.G.K. wrote the manuscript with help and input from all authors.

## STAR-Methods

### Lead contact and materials availability

Further information and requests for resources and reagents should be directed to and will be fulfilled by the Lead Contact, Ilona C. Grunwald Kadow.

### Experimental model and subject details

All experiments were carried out with laboratory-raised *Drosophila melanogaster*. In order to cover both PPL1 and PAM DANs for imaging, we recombined two transgenic driver lines, namely the TH-Gal4 (PPL) driver and the 58E02-Gal4 (PAM) driver, onto the third chromosome. The recombined line was crossed to the UAS-GCaMP6f reporter line for *in vivo* imaging. For experiments using light-field imaging, the additional data to the dataset from [51] was obtained with flies where same TH- and 58E02-Gal4 drivers were recombined again with the UAS-GCaMP6F transgene to obtain a true breeding stock. All fly stocks and crosses were raised on a standard cornmeal medium and maintained at 25°C and 60% humidity in 12 h light and 12 dark cycle climate chambers.

#### Number of experiments

‘N’ indicates the number of flies, while ‘n’ signifies the number of experiments. A total of 319 *in vivo* 2 photon experiments were analyzed from 165 adult female flies of different states stimulated with different odors. (1) Vinegar: fed: n=10, N=8; starved: n=30, N=23; (2) Yeast: fed: n=14, N=11; starved: n=30, N=22; (3) Isoamyl acetate: fed: n=12, N=8; starved: n=13, N=10; (4) 1-Hexanol: fed: n=8, N=4; starved: n=12, N=8; (5) 2-Heptanone: fed: n=9, N=5; starved: n=12, N=8; (6) Ethanol: fed: n=6, N=4; starved: n=7, N=5; (7) 3-Octanol: fed: n=8, N=6; starved: n=9, N=7; (8) 4-MCH: fed: n=11, N=5; starved: n=11, N=9; (9) Peppermint: fed: n=12, N=8; starved: n=13, N=7; (10) Citronella: fed: n=11, N=8; starved: n=38, N=30; (11) Geosmin: fed: n=9, N=5; starved: n=6, N=4; (12) cVA: mated: n=14, N=6; Virgin: n=14, N=6. Odor stimulation was randomized for different experiments and animals.

## Method Details

### Two-photon *in vivo* calcium imaging

4-8 days old female flies were used for all experiments. For starvation experiments, flies of at least 4 days old were transferred to a starvation bottle with only a wet tissue paper in the bottom and a folded wet Whatman round filter paper as a wick hanging from the plug. Starvation was carried out for 24 h or 48 h for the experiments.

For *in vivo* imaging, a fly was restrained in a truncated pipette tip and only a part of the head with antenna and maxillary palp were protruding (see Figure 1A). Proboscis and legs were kept inside the pipette tip to restrain movement. Using fine forceps, cuticle on the dorsal head was removed and the brain was further exposed by removing fat bodies and trachea. The exposed brain was first washed with imaging saline, and then a drop of 1% low temperature melting agarose (NuSieveGTG, Lonza) diluted in imaging saline maintained at 37°C was added on top of the brain in order to minimize brain movement. Imaging saline was added on top once the agarose had hardened. Preparations were imaged with an Olympus 40× 0.8 NA water immersion objective on an Olympus FV1000 two-photon system with a BX61WI microscope. GCaMP6f fluorescence was excited at 910 nm by a mode-locked Ti:Sapphire Mai Tai DeepSee laser. Time series images were acquired at 210 × 210 pixel resolution with 3 frames / s speed using the Olympus FV10-ASW imaging software. For each specimen, first the dorsal and ventral ends of mushroom body DANs were marked in the software. The entire volume of the MB from dorsal to ventral was ~100 μm. Planes were spaced 1 or 2 μm apart in all experiments. Each plane was scanned in time series mode for 80 frames with a speed of 3 frames per second to obtain sufficient spatial resolution for pre-, post- and stimulus phase. On each plane, a 1 sec odor pulse was delivered at the 10^th^ time frame by triggering an automated odor delivery system. After scanning one plane, the focus was shifted to the next plane and the stimulation protocol was repeated for every plane. For each fly, the full volume of the MB was first scanned from dorsal to ventral and then from ventral to dorsal. The type and sequence of odors was randomized over experiments. For instance, in one set of experiments, two different odors were used for the opposite scanning directions, while in another set of experiments, the same odor was used for both scanning directions. In some cases, if the fly was still fit, a third or even fourth odor was recorded.

### Odor stimulation

The following odorants were used in the study: vinegar (Balsamic vinegar, Alnatura, Germany), yeast (Fermipan, Italy; 1g/5ml water), citronella, peppermint (both from Aura Cacia, USA), 3-octanol, ethanol, 4-methylcyclohexanol (MCH), geosmin (0.01% in paraffin oil), isoamyl acetate, 1-Hexanol, 2-Heptanone (all from Sigma-Aldrich, Germany), 11-cis-Vaccenyl acetate (cVA) (Pherobanks, The Netherlands). Odors were diluted in water or paraffin oil to 1% with the exception of yeast and geosmin according to their solubility. A custom-made odor delivery system with mass flow controllers (Natec sensors, Garching) controlled by a MATLAB script were used for odor delivery. Throughout the experiments, a charcoal filtered continuous air stream of 1l/min was delivered through an 8 mm Teflon tube positioned 10 mm away from the fly antenna. Odor was delivered into the main air stream by redirecting 30% of main air flow for 1 s through a head-space glass vial containing 5 ml of diluted odorant. The odor delivery was automatically triggered by counting the time frame output from the scanning microscope by a custom written MATLAB script.

### Image analysis and 3D registration

After acquisition, each imaging plane was first registered in time to correct for within-plane shifts. Then, all the planes were aligned with each other to maintain between-plane continuity. Both registration steps were performed with standard methods implemented in the scikit-image Python package (register_translation) [53].

Due to the variability in GCaMP expression between MB compartments and flies and lack of significant landmarks in the region imaged, automatic registration was not possible. Therefore, to assign activity to lobes, we used a semi manual-procedure. The binary masks for lobes published by Aso et al. [11] were then converted to meshes and simplified with the vtk toolkit. Next, the Aso et al. reference MB was registered manually to a high-resolution reference volume of the line TH-Gal4/58E02-Gal4;UAS-GCaMP6f used in this study (Fig. S1.2, step 1). Since the mushroom body is a L-shaped structure, determining the location of three points in three dimensions is enough to specify the location, rotation and scale while also allowing for bending between the α, and β and γ lobes. Using a custom graphical user interface tool three points were determined: at the ends of the lobes, and the point where α and γ lobes come together. The exact location of the registration point is not important for the alignment quality, as long as they are located at the same cell, axon bundle or other landmark feature in both stacks.

Once this correspondence was established, the same procedure was repeated between each imaging experiment and the high-resolution TH/58E02 reference (Fig. S1.2, step 2). Finally, for each transformed lobe or compartment, time traces of all voxels belonging to it (determined with a ray-intersection method from the trimesh library (developed by Dawson-Haggerty et al. version 3.2.0 at https://trimsh.org/) were extracted and saved per plane (each plane being a separate repetition, Fig. S1.2, step 3).

The developed tool provides several advantages over the Longair and Jefferies ImageJ plugin (https://imagej.net/Name_Landmarks_and_Register): it shows an interactive preview of the segmentation, works with mesh-, instead of voxel-defined regions and is significantly faster to use. The ease of use was an important issue given the several hundreds of stacks acquired through the experiments.

For imaging experiments on the light field microscope, landmark registration was performed with FIJI’s landmark_registration plugin (http://imagej.net/Name_Landmarks_and_Register) with the template of MB compartments.

### *In vivo* light field imaging

Light field imaging was carried out as previously described [51]. Briefly, the experiment to obtain data with a walking fly was performed on a custom-built light-field microscope using a microlens array to separate rays coming from different angles [54]. The resulting light field images have information about sample depth, and deconvolution allowing to reconstruct the volume. This further allows to record the whole volume without scanning. Our light field microscope was constituted of a Leica HC FLUOTAR L 25×/0.95 objective and an MLA-S125-f10 microlens array from RPC photonics in a Thorlabs Cerna system. The microlens array was placed on the image plane, while the camera imaged the microlens array through 50 mm f/1.4 NIKKOR-S Nikon relay lenses. The light field images were recorded at 25Hz with a scientific CMOS camera (Hamamatsu ORCA-Flash 4.0). The volumes were reconstructed offline, using a python program developed by [54] and available on github: https://github.com/sophie63/FlyLFM.

### Quantification and Statistical Analysis

#### Statistics and other types of data analysis

To reduce noise, we averaged responses during four time points after odor onset. Since for the majority of recorded flies, only two odors per fly were recorded or analyzed, we suspected strong multicollinearity in the data. Indeed, variance inflating factors were high and sometimes infinite. After removing the fly identity factor, variance inflating factors were three or less (without considering interactions), indicating low multicollinearity. We thus removed fly identity from the analysis (however as described below, we performed a mixed model to verify that fly identity did not strongly affect regression coefficients).

We next performed ANOVA, using a model considering all remaining factors (stimulus, position in odor sequence, imaging direction, starvation state), as well as compartment-specific effects (Table 1). As variance explained by position in odor sequence (i.e. whether odor was applied first or second in the sequence) was small and non-significant, we removed this factor for further analysis. We used this model for estimating regression coefficients:

> “df_f ~ lobe*(stimulus+order_presented+dorsal_to_ventral_val)+starved:lobe-1“.

As cross-validation indicated some overfitting (24 % instead of 37 % of variance explained out of sample data), we also performed an elastic net regularization. We also used a mixed model with fly identity as group, to account for any fly specific effects. For coefficients related to valence, the same models were used but replacing stimulus with valence, and restricting the data to odors with clear valence (vinegar, yeast, citronella, geosmin, and peppermint). Finally, we estimated coefficients using minimal models for lobe (“df_f ~ lobe-1”), starvation (“df_f ~ lobe+lobe:starved-1”), and valence (“df_f ~ lobe+lobe:valence-1”). The coefficients for all the models are in good agreement (Fig. S1, S3 and S4). As cVA was the only odor recorded in the presence of different mating states, we used a simple model (“df_f ~ lobe+lobe:virgin_val-1”) on this data subset to evaluate coefficients related to mating. We also performed post-hoc t-tests for lobe specific effects of starvation, valence, and mating. All the p-values were pooled and corrected for multiple comparison using a Simes-Hochberg procedure. For walking experiments, we only performed regression of lobe averaged activity with behavior time series convolved with a GCaMP6 kernel and normalized to have an average value of one during walk (to be able to compare regression coefficients with other factors with 0 or 1 values). The ANOVA and regression analysis were performed using statsmodels (http://statsmodels.sourceforge.net/).

To see whether the fly can theoretically decode compartment activity to get information about stimulus, valence or starvation state, we trained logistic regression classifiers (lbfgs solvers for all, and balance class weight and multinomial logistic model for stimulus decoding). We also projected the data to obtain one to three dimensions using linear discriminant analysis. We used the three first loadings to construct the odor space in Fig. 2. The LDA and classification were performed using scikit-learn [55].

## Supplementary Figures

**Figure S1.**
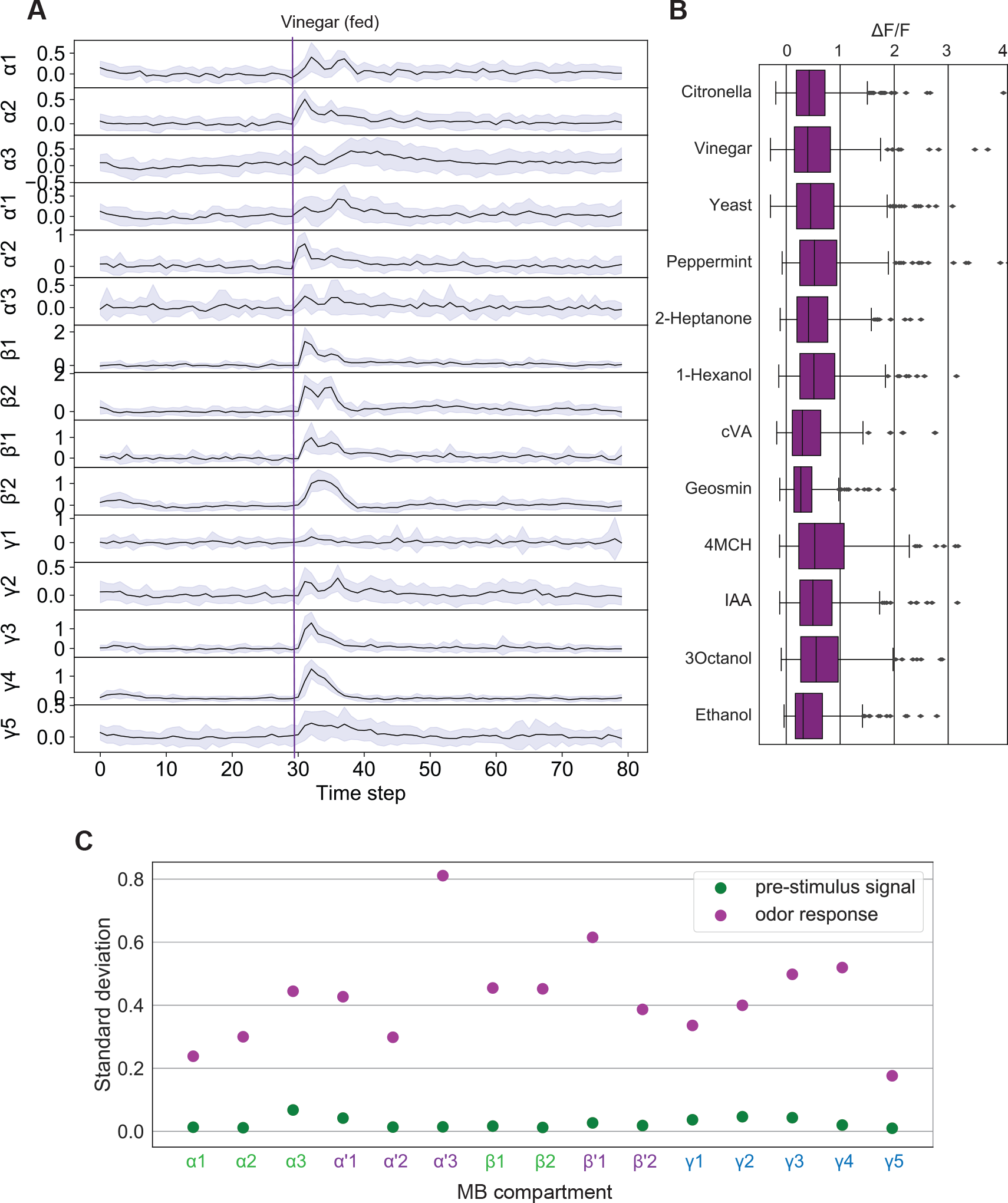
Related to Fig. 1. **(A)** Example of average (and 95% interval) of individual planes time series (corresponding to individual odor presentation) for each compartment. **(B)** Odor specific responses. The data for all lobes was pooled. **(C)** Odor response standard deviation for data shown in (H, pink) and standard deviation of the pre-stimulus phase (I, green).

**Figure S1.2.**
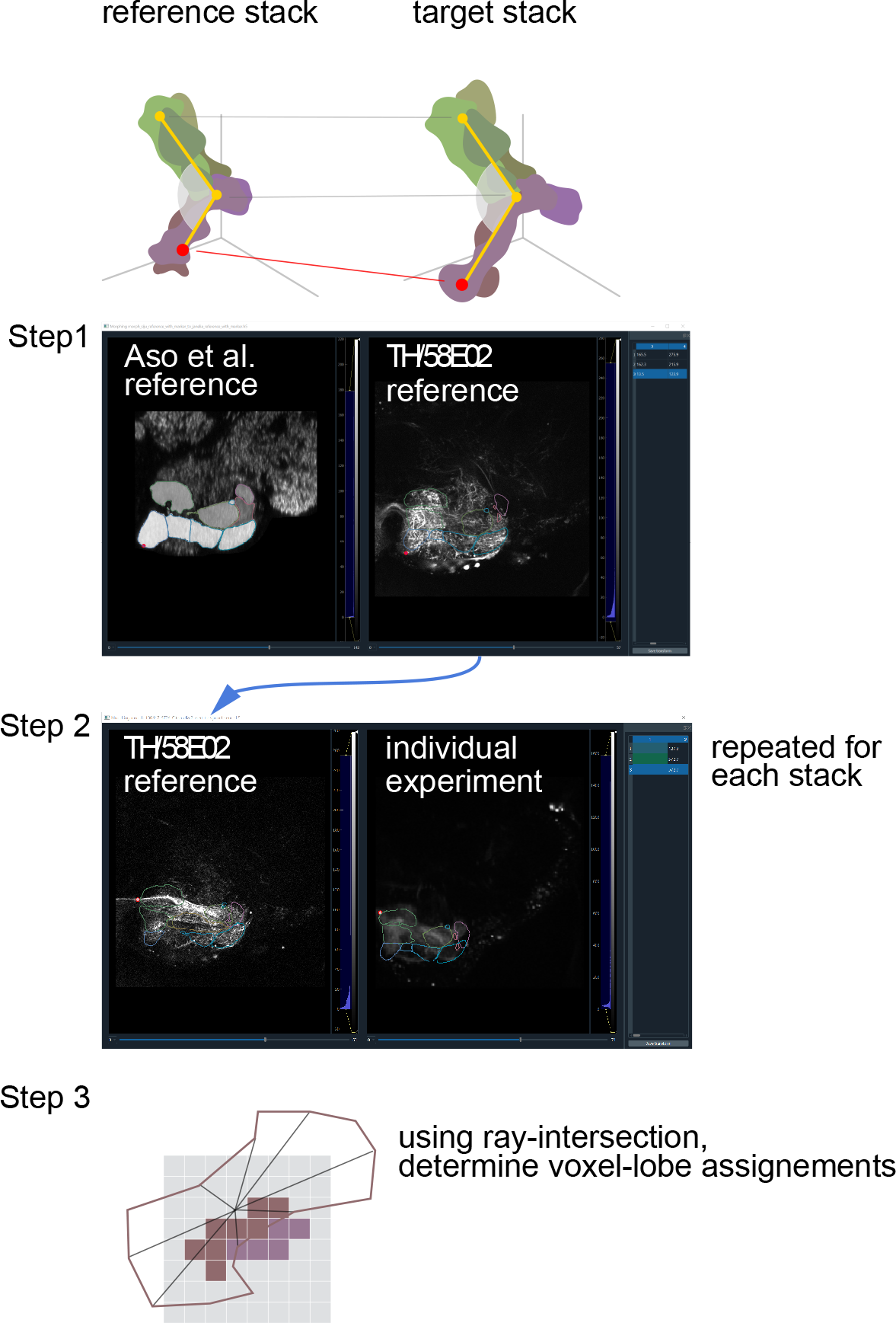
Related to Fig. 1. Lobe alignment procedure: three points are used to place lobe meshes into the coordinate system of an individual experiment. The points are matched first across the Aso et al reference, and a high-resolution stack of the imaged line (step 1), and then the individual, lower-resolution experiments, to the high-resolution version of the same line (step 2). The meshes transformed in this way are used to assign voxels to lobes (step 3).

**Figure S2.**
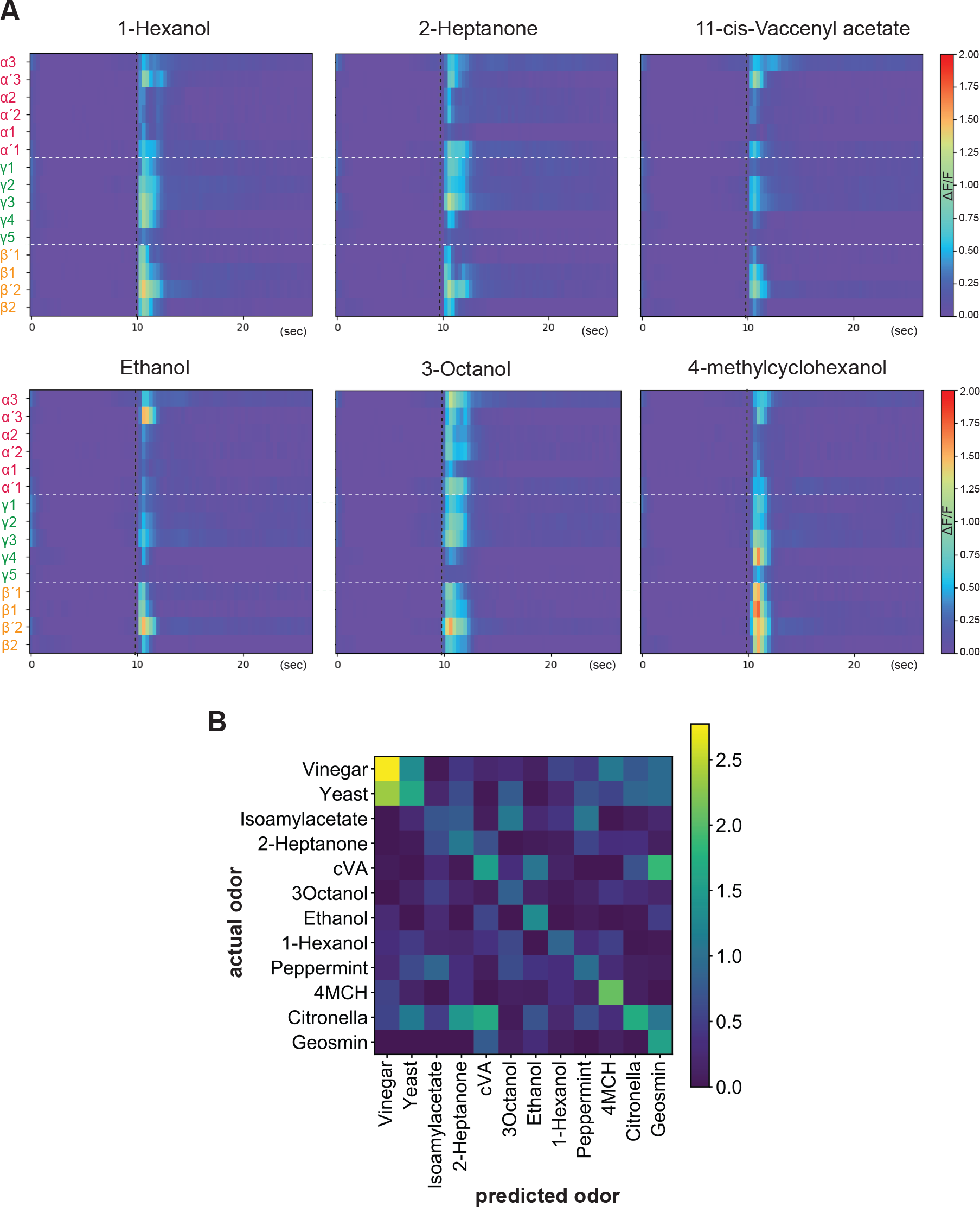
Related to Fig. 2. **(A)** Heatmaps showing averaged DAN responses to different odors in 15 mushroom body compartments (n = 165 female flies, all states). Vertical dashed black line indicates stimulus delivery time. Horizontal dashed white line separates group of lobes. **(B)** Confusion matrix for the SVM classification showing how well odor identity was predicted from the analysis of the DAN population data.

**Figure S3.**
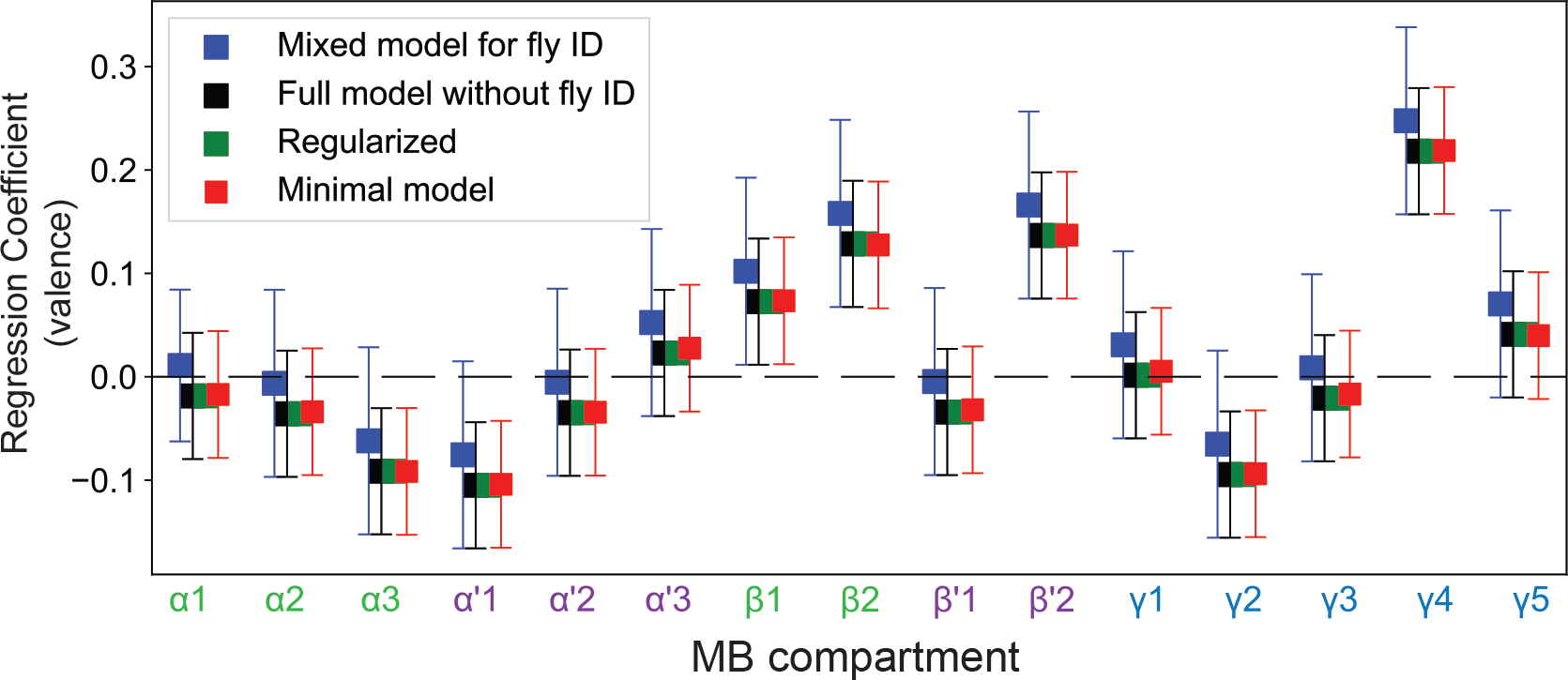
Related to Fig. 3. Valence regression coefficient for specific mushroom body compartments. Details of the four different models are described in the method section.

**Figure S4.**
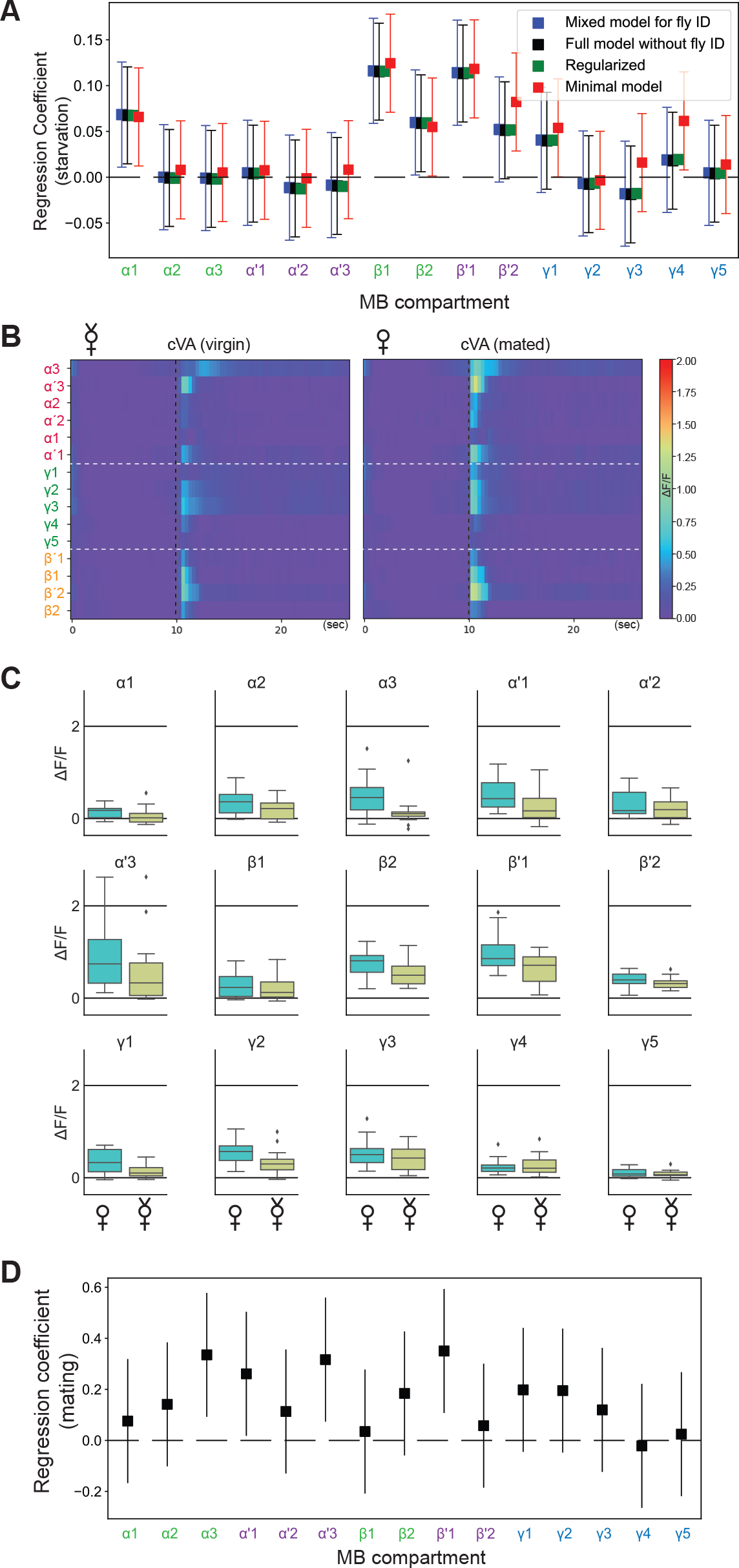
Related to Fig. 4. **(A)** Starvation regression coefficients for specific lobes. Model details are described in the method section. **(B)** Heatmaps showing averaged DAN responses to 11-cis-Vaccenyl acetate (cVA) from virgin (n = 14) and mated females (n = 14). Vertical dashed black line indicates stimulus delivery time. Horizontal dashed white line separates group of lobes. **(C)** Pairwise comparisons of mated vs. virgin female flies for MB compartment-specific responses to cVA. **(D)** Mating state regression coefficients for specific MB compartments.

**Figure S5.**
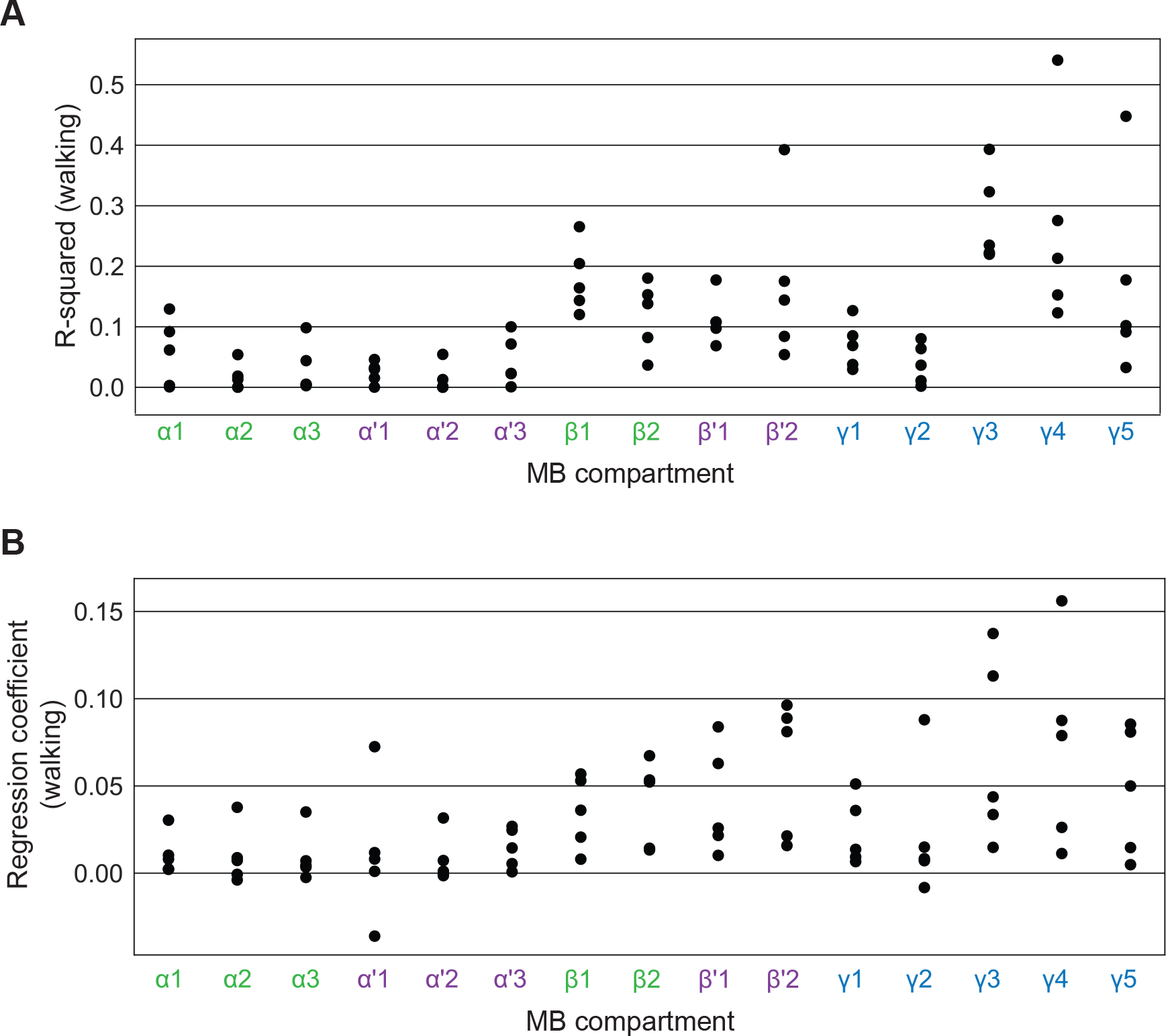
Related to Fig. 5. **(A)** R-squared of regression with walking in light field experiments in which no odor was presented to a fly walking on a ball. N = 5 flies. **(B)** Coefficients for regression with walking. N = 5 flies.

