## Supplementary material for "Valence and state-dependent population coding in dopaminergic neurons in the fly mushroom body": Table 1

**Table 1. ANOVA table showing the variance associated with different factors**

| | **sum of squares** | **% of variance explained** | **PR(>F)** |
|---|---|---|---|
| **MB compartment** | **279.9** | **23.2** | **3.00E-302** |
| **stimulus** | **40.1** | **3.3** | **6.60E-45** |
| **Compartment : stimulus** | **106.5** | **8.8** | **9.30E-57** |
| order presented | 0.6 | 0.1 | 5.30E-02 |
| Compartment : order_presented | 2.5 | 0.2 | 3.80E-01 |
| imaging direction | 2.7 | 0.2 | 4.70E-05 |
| Compartment : imaging direction | 6.2 | 0.5 | 5.40E-04 |
| starvation state | 2.7 | 0.2 | 5.70E-05 |
| Compartment : starvation state | 6.2 | 0.5 | 5.70E-04 |
| Residual | 760.6 | 63 | |

**$R^2$ = 0.37383763806006964**

* Most significant factors are in bold.
