## Supplementary material for "Valence and state-dependent population coding in dopaminergic neurons in the fly mushroom body": Table 2

**Table 2. ANOVA table showing variance associated with valence**

| | sum of squares | % of variance explained | | PR(>F) |
|---|---|---|---|---|
| **MB Compartment** | **174.5** | **26.6** | **26.6** | **1.5E-174** |
| Valence | 0.4 | 0.1 | **3.6** | 1.2E-01 |
| **Compartment : Valence** | **21.4** | **3.3** | | **2.6E-19** |
| Imaging direction | 0.9 | 0.1 | 0.8 | 2.1E-02 |
| Compartment : Imaging direction | 4.5 | 0.7 | | 2.4E-02 |
| Position in odor sequence | 0.5 | 0.1 | 0.5 | 9.4E-02 |
| Compartment : Position in odor sequence | 2.7 | 0.4 | | 3.5E-01 |
| Starvation | 2.4 | 0.4 | 0.8 | 1.9E-04 |
| Compartment : Starvation | 2.7 | 0.4 | | 3.3E-01 |
| Residual variance | 445.2 | 68.0 | 68.0 | |

**$R^2$ = 0.32**

**Adj. $R^2$ = 0.30**

* Most significant factors are in bold.
